# Use of Solentim verified in-situ plate seeding (VIPS™) enhances single-cell cloning efficiency

**DOI:** 10.1101/2022.06.03.494661

**Authors:** Nadia Ilyayev, Catherine Martel, Mostafizur Mazumder, Nishi Singh, Ramtin Rahbar

## Abstract

The primary goal in cell line development is to establish high-producer recombinant cell line(s) of single-cell origin. Traditionally, these cell lines are developed using limiting dilution cloning (LDC) method of single cell isolation, a rate-limiting, lengthy and labor-intensive process.

The Verified-In-Situ-Plate-Seeding (VIPS™) is an automated single-cell seeding and imaging equipment designed to accelerate cell line development workflow. In this study, VIPS™ was tested for efficiency and accuracy of single cell seeding in parallel with limiting dilution cloning (LDC). Three Chinese hamster ovary (CHO) derived cell lines with known clonal properties were tested under six different growth conditions (three growth media and two different kinds of microplates). Data showed VIPS™ and limiting dilution (LDC) have comparable cloning efficiency when CHO-M cells were tested. By contrast, the Verified In-Situ Plate Seeding (VIPS™) produced 6-8-fold more clones of single-cell origin than LDC when CHO-K1 or CHO-S cells were tested. Moreover, the verified In-Situ Plate Seeding (VIPS™) correctly identified single-cell and multiple-cells seeded wells with 65-72% and 52-81% accuracy, respectively. Taken together, the high throughput imaging and single-cell seeding capabilities of VIPS™ outperformed the rate-limiting LDC method and, therefore, has the potential to accelerate cell line development workflow.

## Introduction

Mammalian Cell Culture (MCC) is a popular platform for manufacturing of biopharmaceuticals such as monoclonal antibodies (mAbs), Fc-fusion and other recombinant proteins (1).

International Conference on Harmonization (ICH) of Technical Requirements for Registration of Pharmaceuticals for Human Use (2), as well as the United States Food and Drug Administration (FDA) (3) stipulate that for production of biopharmaceuticals, the cell substrate expressing the desired product must be of pure clonal origin.

Isolation of single cells for the purpose of establishing clonally pure cell lines is usually performed using the Poisson distribution analysis-LDC method (4). In LDC, a mixed population is serially diluted and seeded into 96-well plates, such that each well receives no more than a single cell (4). The clonal purity of cell lines established by a single round of LDC has been questioned due to the statistical nature of the method and cellular tendency to form aggregates. Therefore, it is recommended that two consecutive rounds of LDC be performed to ensure clonality of the resulting cell line(s) (5, 6). However, even after two consecutive rounds of cloning at 0.3 cell/well density, only 75% of the clones can be considered monoclonal at 95% confidence, as estimated by Poisson statistical analysis (4). Although the probability of clonality may be further increased by using lower seeding density, the number of cell lines recovered will be reduced (4). Due to the rate-limiting nature of the single cell clone isolation, LDC is ultimately an inefficient tool for building a large pool of clonal cell lines for use in screening. To address the issues of low cloning efficiency and absence of direct and reliable proof of clonality, Solentim introduced the VIPS™ system, a high-capacity single-cell seeding and imaging equipment (7) and Cell Metric®, an imaging system to follow clonal growth from seeding to full growth (8).

VIPS™ dispenses cells in nanoliter size droplets while its z-stacking capable imaging system scans for the presence of cells in each droplet, as each droplet is deposited. At the first instance of cell detection, VIPS™ terminates further seeding to ensure no additional cells are seeded in the same well. VIPS™ image-analysis-algorithm verifies the z-stack images and labels each well accordingly as single cell, more than 1 cell, and null cells in real-time. A case study by Jensen and Jensen demonstrated that VIPS™ seeds verified single cells in 87% of the wells in a 96-well microplate (7).

Cell Metric® is a high-resolution imaging system specifically designed for monitoring clonal growth and verification of clonality (8). The image analysis software accurately and unequivocally identifies single cell derived clones. The automatic focus system measures and adjusts the optimal focus in each well and uses a precision XYZ positioning system to eliminate blurring due to “image stitching”. The automatic focus system results in clear visualization of 100% of the wells with uniform imaging up to and including the well edge. This image-based approach provides a permanent record of evidence in three ways: i) a clear image of each well starting at Day 0 (day of plating), ii) images of wells during the growth period, and iii) an objectively measured growth rate for each colony. An audit trail for the growth of a single cell can then be provided with photo-documentation. When paired together, VIPS™ and Cell Metric®, are expected to form a powerful functional unit which not only has the potential to increase cloning efficiency but also allow tracking of complete clone history from seeding to full clonal growth. For these reasons, both equipment have been adopted globally with unit installations at different sites.

In this study, using three different CHO cell lines we proceeded to compare the single cell cloning efficiency of VIPS™ with that of the LDC under six different growth conditions (three growth media, and two different tissue culture microplates). Our data suggest that, for developing cell lines, VIPS™ has the capability to produce 6 – 8-fold more single-cell derived clones than the traditional LDC method.

## Materials and Methods

### Cell lines

Three CHO cell lines, code named CHO-1, CHO-2, and CHO-3 were used in the study. CHO-1 is a proprietary CHO-M cell line expressing a humanized IgG4 mAb; CHO-2, a CHO-K1 cell line, were developed in-house and express EGFP and a puromycin resistance gene; CHO-3 is a naïve proprietary CHO-S cell line.

### Cell culture conditions

The cells were grown in 125 mL polycarbonate baffled shake flasks (VWR, Catalogue # 89095-258) at an agitation speed of 120 revolutions per minute (rpm) at 37°C, 80% relative humidity (RH) and 8% CO_2_. Cells were passaged every 3^rd^or 4^th^day by seeding in fresh growth medium at 0.3 × 10^6^cells/mL.

### Growth medium, Conditioned Medium (CM) and Cloning medium

CM was obtained for each cell line by growing the cells in their corresponding growth medium (Fig 1 and Table 1) and collecting the supernatant of the cultures in the logarithmic (log) growth phase. The supernatants were collected by centrifugation (300 x g, 5 min, room temperature) and filter-sterilized using a 0.22 µm filter (Millipore, Catalogue # SLGP033RS).

**Table 1:** List of media and supplements. Three growth media, one cloning medium, and supplements used for seeding and cultivating three different CHO cell lines.

| Media/Supplement | Vendor | Catalogue# |
| --- | --- | --- |
| ExCell Cloning medium | Sigma | C6366 |
| HT supplement | Life Technologies | 11067030 |
| Glutamine | Life Technologies | 35050061 |
| SFM4CHO | GE Healthcare | SH30548.02 |
| ExpiCHO Expression Medium | Life Technologies | A2910003 |
| ExCell Advanced Medium | Sigma | 14366C-1000ML |
| Cell Boost 5 | GE Healthcare | SH30865.02 |

**Fig 1.**
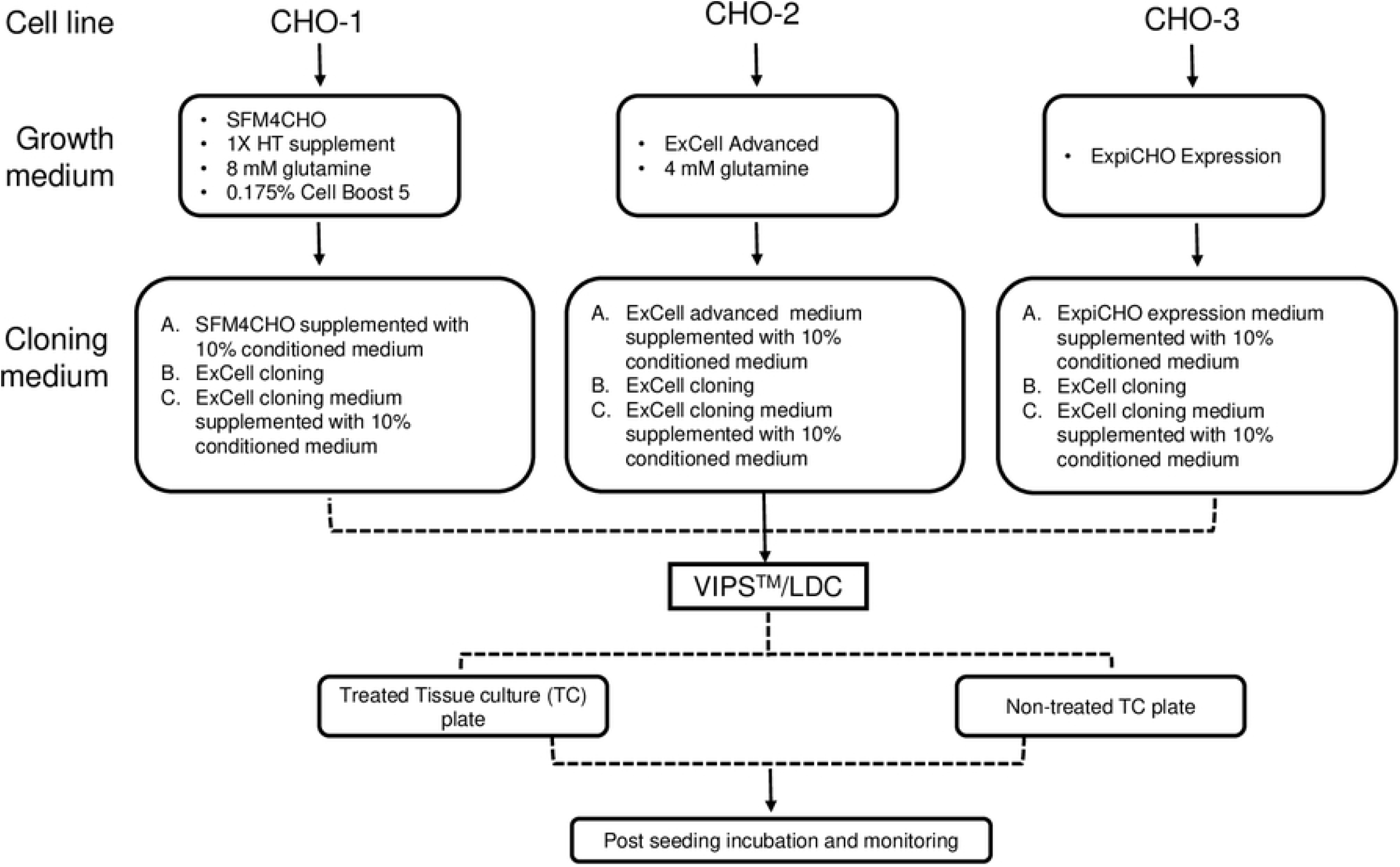
Cell seeding workflow. Three different cell lines (CHO-1, CHO-2, and CHO-3) were grown in their corresponding growth medium to produce conditioned medium (CM) (see materials and methods). Treated tissue culture (TC) and non-treated TC microplates (96 well) were used for seeding in parallel for both VIPS™and LDC experiment using cloning media A, B and C. Seeded plates were incubated at 37°C with 8 % CO_2_ and cell growth was followed for 28 days after seeding.

### Preparation of cell suspension for seeding

For LDC, cell cultures were diluted by serial dilution to achieve a final concentration of 5 cells/mL in the appropriate cloning medium (Sigma, Catalogue # C6366). 200 µL of diluted cells were dispensed into each well of treated tissue culture (TC) and non-treated TC 96-well microplates to obtain an average seeding density of 1 cell/well. For VIPS™, a 10,000 cells/mL suspension prepared in cloning medium was used for seeding the 96-well microplates.

### Tissue culture plates

Treated tissue culture (TC) (Corning, Catalogue # 3598) and non-treated tissue culture (TC) (VWR, Catalogue # 10861-562) plates were used.

### Plate incubation and monitoring

After seeding, the plates were centrifuged at 300 x g for 2 min and scanned using the Solentim Cell Metric® instrument. Cells were incubated at 37°C, 8% CO_2_ and 80% RH. Culture plates were scanned on day 0 (seeding day), day 1, day 2, day 3, day 7, day 14, day 21 and day 28. Wells showing cell growth were examined using Cell Metric® software to determine clonality.

#### Experiment 1

Three cloning media were tested for each of the three cell lines. The tested media were: (A) cell-specific growth medium supplemented with 10% CM, (B) ExCell cloning medium, (C) ExCell cloning medium supplemented with 10% CM. The CHO-2 cell line growth medium contained 7.5 µg/mL puromycin (Thermo Fisher, Catalogue # A1113802) and each of the cloning media tested for CHO-2 were supplemented with 7.5 µg/mL puromycin accordingly.

#### Experiment 2

Cloning efficiency of VIPS™ was compared with that of manual LDC using CHO-1 cell line grown in Medium C.

## Results and Discussion

### Single cell seeding using VIPS™ is significantly more efficient than LDC

To compare cloning efficiency of VIPS™ and LDC, a series of small-scale seeding efficiency assays were executed using three different cell lines (CHO-1, CHO-2, and CHO-3), each seeded in three different cloning media (Medium A, Medium B, and Medium C). Cells were seeded in a 96-well plate at 1 cell/well by LDC or VIPS™. Cell seeding workflow is shown in Fig 1.

Fig 2 shows the proportion of wells for VIPS™ and LDC seeded plates combined. Cell growth in cloning medium C for all three cell lines (Fig 2) was observed while and no growth was detected in cloning media A and B. Note that each cell line grew to different proportions in medium C reflecting specific differences in the outgrowth potential for each cell line.

**Fig 2.**
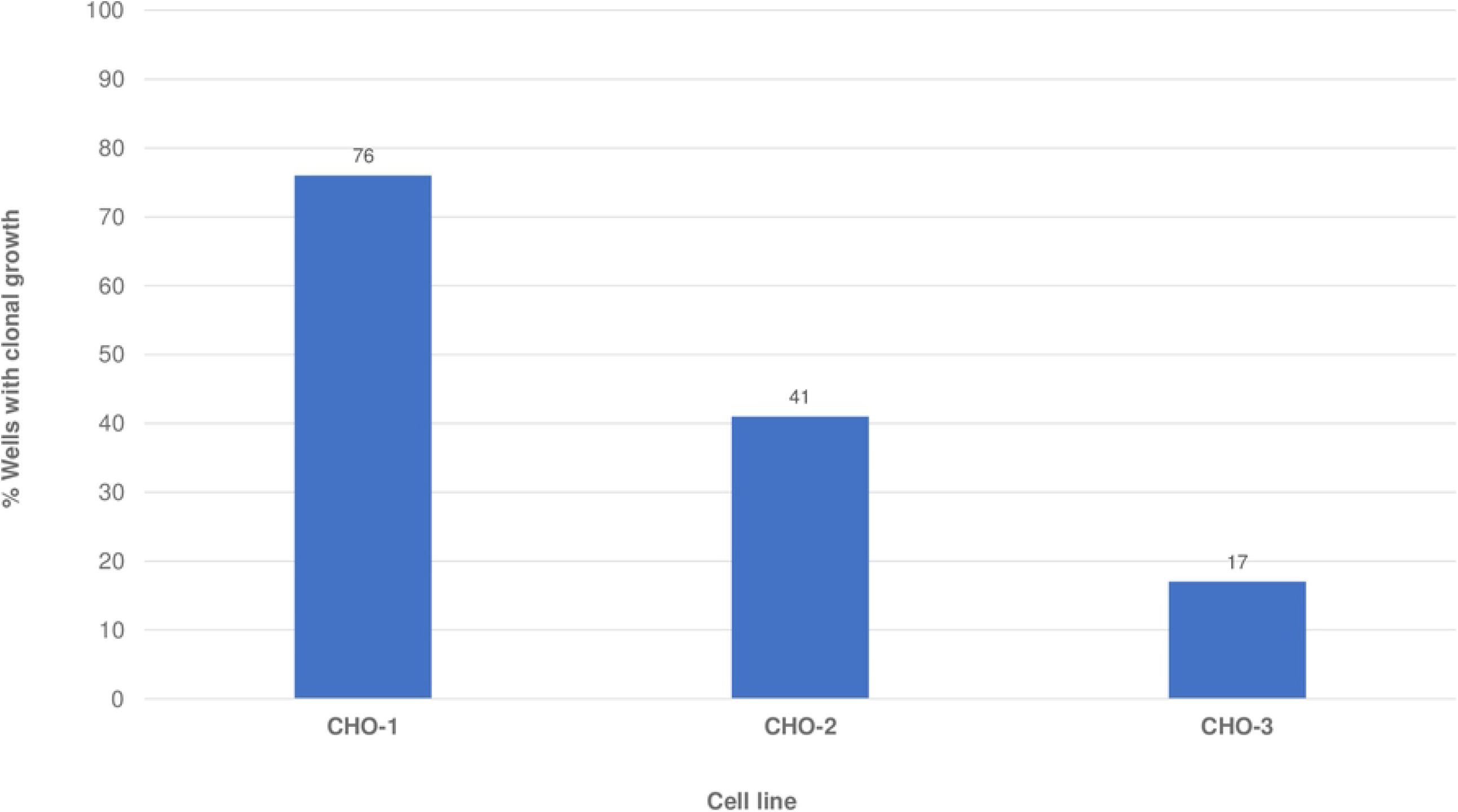
Clonal growth differential. Proportion of wells showing growth of CHO-1, CHO-2, and CHO-3 cells in the cloning medium C.

The number of clonal and non-clonal populations recovered from each seeding method in Experiment 1 and Experiment 2 were compared. VIPS™-seeded wells with CHO-2 and CHO-3 cell lines showed a higher percentage of growth compared to LDC seeded wells. Notably, CHO-1 cell line showed similar growth rate in both Experiment 1 and 2 (Fig 3). Clonality was determined using Cell Metric® whole well imaging from day 0 onward. Seeding using VIPS™ resulted in a 1.2-to-7.8-fold improvement in the number of clonal populations recovered for the three cell lines (Table 2).

**Table 2:**
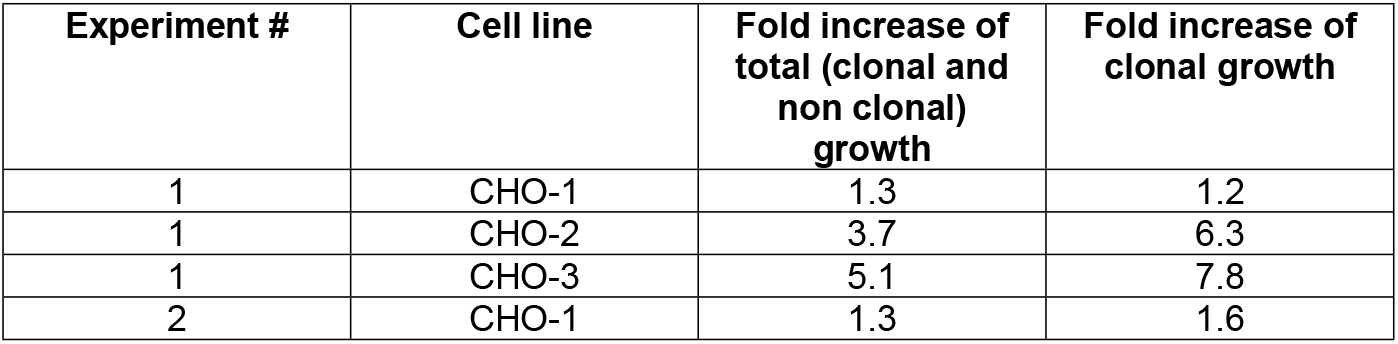
Fold increase in growth of cells (clonal as well as non clonal) seeded using VIPS™.

| Experiment # | Cell line | Fold increase of total (clonal and non clonal) growth | Fold increase of clonal growth |
| --- | --- | --- | --- |
| 1 | CHO-1 | 1.3 | 1.2 |
| 1 | CHO-2 | 3.7 | 6.3 |
| 1 | CHO-3 | 5.1 | 7.8 |
| 2 | CHO-1 | 1.3 | 1.6 |

**Fig 3.**
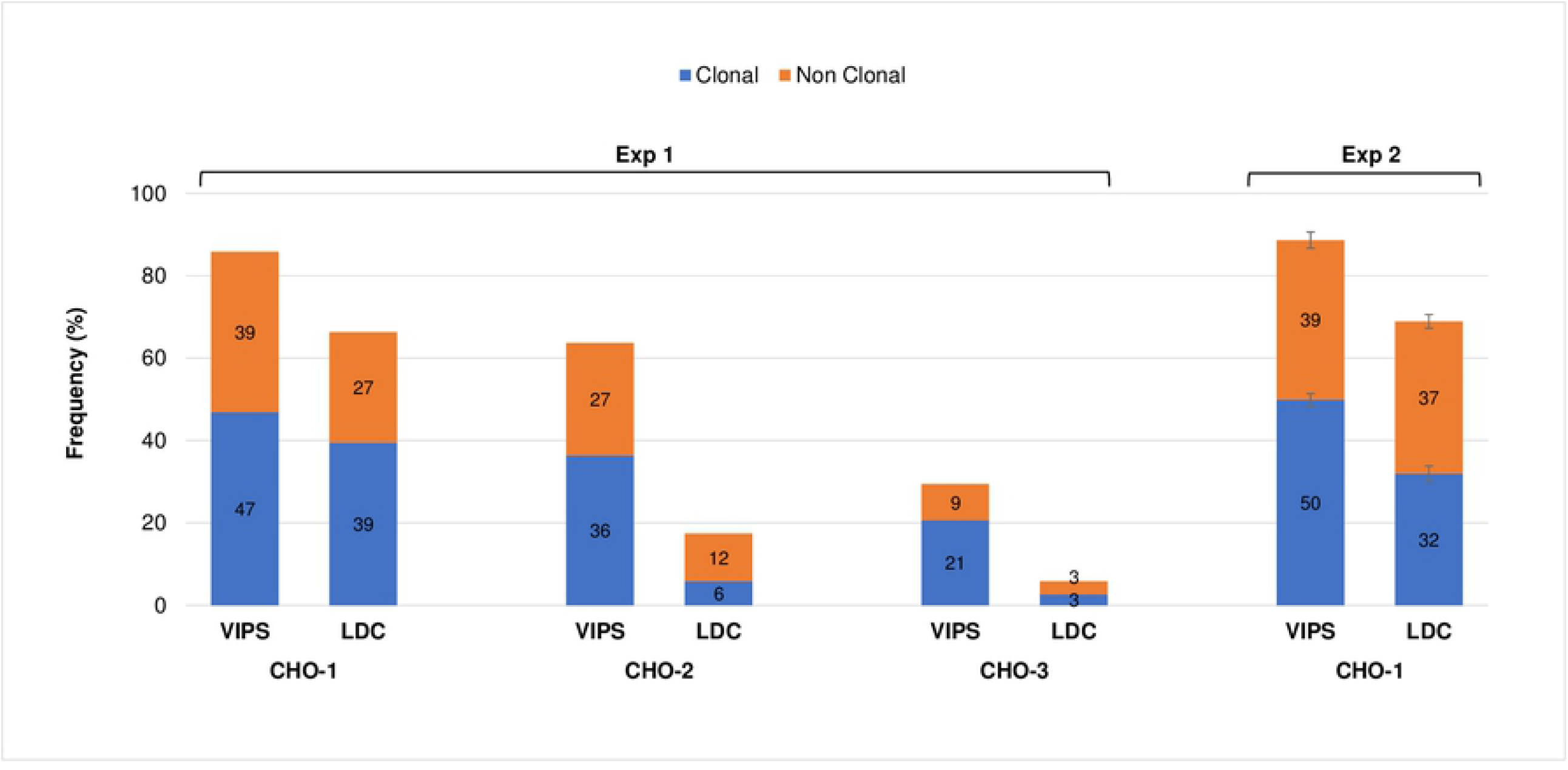
Clonality Distribution of plates seeded using VIPS™ and LDC at 1 cell/well. Experiment 1 consists of 2 plates per condition. Experiment 2 consists of CHO-1 cells seeded on 5 plates per method. Error bars for Experiment 2 represent standard error.

The use of VIPS™ had a more pronounced effect on the cloning efficiency of CHO-2 and CHO-3 lines with a 6.3- and 7.8-fold increase, respectively (Table 2). Of note, the distribution of clonal vs non-clonal wells in Experiment 1 CHO-1 LDC-seeded plates is in agreement with the expected Poisson Distribution for a 1 cell/well seeding density (Table 3).

**Table 3:** Poisson distribution of cells in LDC.

| Cell/well | Frequency |
| --- | --- |
| 0 | 37% |
| 1 | 37% |
| ≥2 | 26% |

This suggests that CHO-1 cell line has an outgrowth potential close to 100% in cloning medium C indicating that almost all cells seeded were able to grow. VIPS™ CHO-1 clonality distribution observed in Experiment 2 was similar to the observation made in Experiment 1; 50% of the wells on average contained clonal populations. In contrast, LDC-seeded plates in Experiment 2 had more non-clonal populations (37%) and less clonal populations (32%) than what is expected (37%) from the Poisson distribution (Table 3). The results indicate the inefficiency inherent to LDC method and the advantage and efficiency of the VIPS™ method of single cell seeding.

Fig 4A shows the number of times VIPS™ recorded single-cell, multiple-cell and null-cell seeding events. Across experiments and conditions, VIPS™ showed a single cell seeding efficiency ranging between 75% to 80% (overall average of 77%). CHO-3 cell line showed the highest efficiency with 80% single seeding event recorded. Notably, CHO-3 also had the highest proportion of empty wells (10%) compared to other cell lines which suggest that the initial cell mixture loaded on the instrument was more dilute compared to the other tested cell lines. The results indicate that seeding efficiency can be improved by optimizing the cell concentration loaded onto the VIPS™.

**Fig 4.**
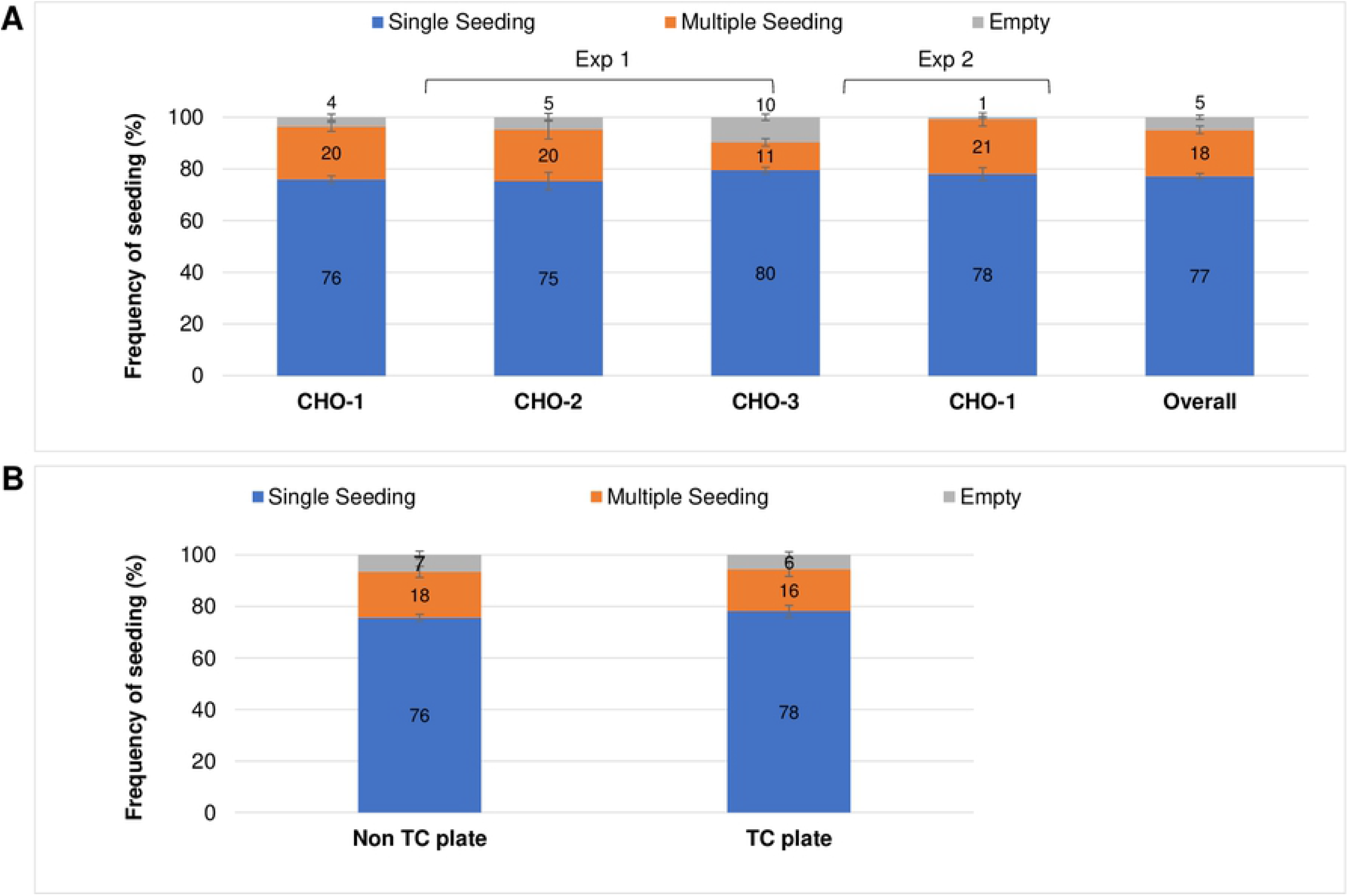
VIPS™ Seeding Efficiency. A) Average frequency of single and multiple seeding events using different cell lines. B) Effect of plate type on seeding efficiency. Error bars in A and B represent the standard error. TC plate = Treated Tissue Culture (TC) plate, non-TC plate = Non-treated TC plate.

### VIPS™ seeding efficiency was similar in both treated TC and non-treated TC plates

To evaluate if VIPS™ has performance bias towards a specific plate type, (a) treated TC, and (b) non-treated TC plates were tested for seeding efficiency. The polystyrene plastic of treated TC plates is modified to reduce its hydrophobicity, promoting cell attachment. Note that the three tested CHO cell lines have been adapted to grow in suspension and do not require treatment of TC for growth. Using treated TC plates provides the opportunity for the seeded cell and their progeny to remain in proximity to each other thus making the clonality assessment easier when using high-resolution imaging instruments, such as Solentim Cell Metric®. Moreover, VIPS™ software is optimized for treated TC plates. Due to the hydrophobicity of the plate type the shape of the nanodroplet deposited by VIPS™ can vary. Nanodroplets deposited on treated TC plates are wider and flatter, whereas nanodroplets deposited on non-treated TC plates are rounder and taller. As shown in Fig 4B, VIPS™ seeded both plate types at a similar efficiency.

### VIPS’ ability to seed single cell in each well was 65% to 72% accurate when verified with the actual growth and clonality report from Cell Metric®

The accuracy of VIPS™ in identifying single and multiple seeding events was tested by examining wells showing growth using Cell Metric® instrument. For clonality assessment, only wells showing cell growth were analyzed and wells with no growth were not included in the analysis. Based on Cell Metric® growth tracking profile of each well, VIPS™ correctly identified single cell seeding events between 65% and 72% of the time and multiple-seeding events between 58% and 81% of the time (Fig 5).

**Fig 5:**
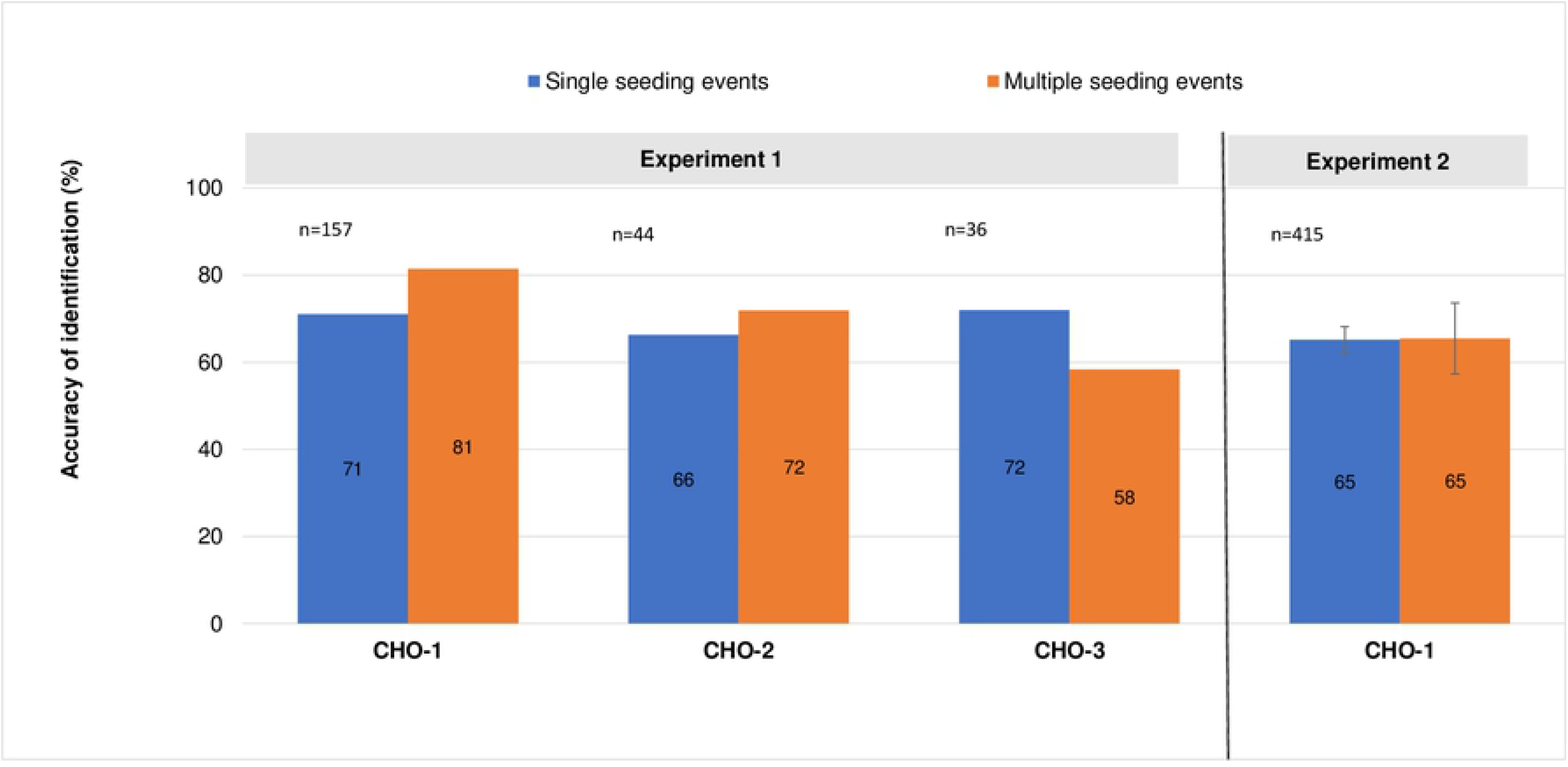
Accuracy of clone identification. Accuracy in clone identification, shown as percent of the total number of clones and “n” represents number of wells with growth.

### Cell Metric® confirmed the growth profile and clonality

As described in materials and methods section, VIPS™ seeded plates were imaged overtime to track the growth profile and to produce FDA-acceptable proof of clonality reports. A Clonality assessment ensures that the growth of a particular cell line emerged from a single cell. Fig 6A shows representative images following the growth of a single cell for 10 days in culture. Fig 6B represents whole well images on Day 1 and Day 10

**Fig 6.**
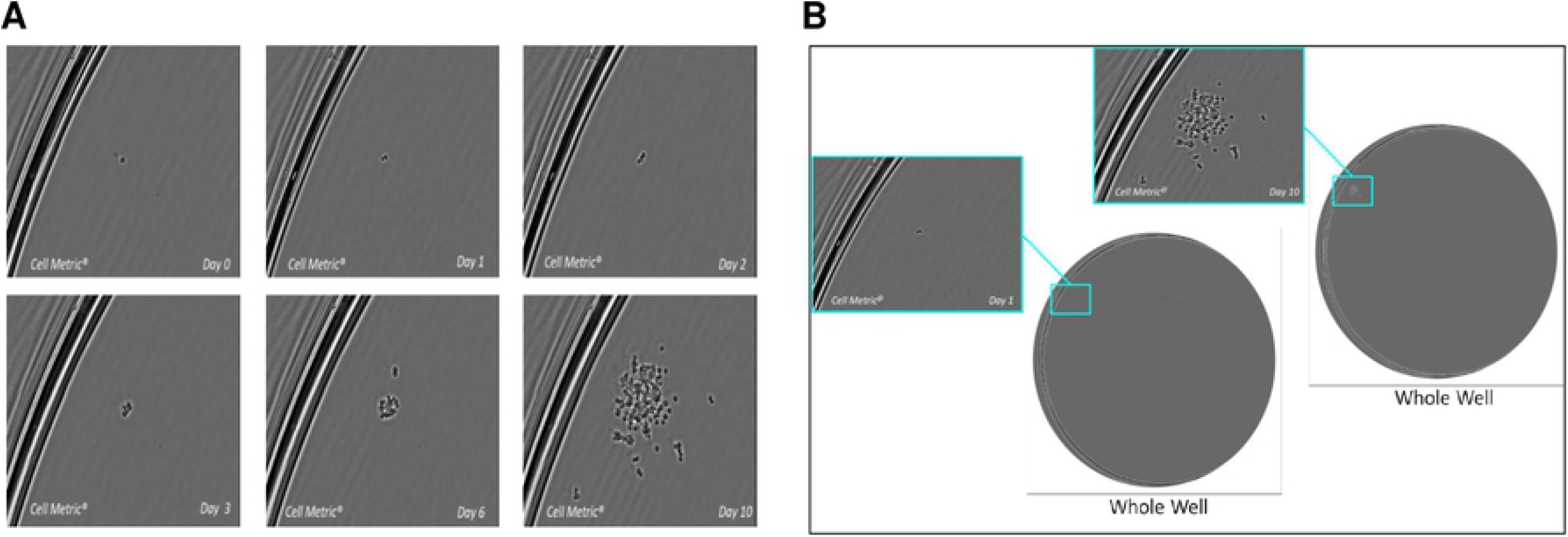
Proof of monoclonality. Images of a single cell outgrowing as a clone on Day 0, 1, 2, 3, 6 and 10 as tracked by Cell Metric® imaging system (A), and whole-well images of the same clone from Day 1 and Day 10 of culturing to demonstrate monoclonality (B).

## Conclusion

Our results demonstrated an improved cloning efficiency of the three cell lines tested using VIPS™ compared to LDC. The improvement was found to be most pronounced for the CHO-K1 and CHO-cell lines with up to 8 folds more clones recovered. Moreover, we showed that a definitive proof of clonality that meets regulatory requirement can be established by using the Cell Metric® platform without having to rely on statistical probability.

## Acknowledgement

No potential conflicts of interest were disclosed. The authors thank Michelle Ng for critically reading the article.

